# Accelerated Aging Signatures in 3D Genome Organization and Transcriptome in Schizophrenia

**DOI:** 10.64898/2026.05.25.727571

**Authors:** K.A. Ulianov, D.R. Zagirova, A.D. Kononkova, A.V. Dudkovskaia, M.N. Molodova, K.V. Morozov, O.I. Efimova, M. Bazarevich, A. Cherkasov, P.D. Morozova, A.V. Tvorogova, I.A. Pletenev, N. Kondratyev, V.E. Golimbet, S.V. Razin, P. Khaitovich, S.V. Ulianov, E.E. Khrameeva

## Abstract

Schizophrenia is a severe neuropsychiatric disorder that affects the behavioral, emotional and cognitive state of patients. Despite its substantial heritability, the molecular etiology of the disease remains poorly understood. Many schizophrenia-associated genetic variants reside in non-coding regions, and exert their effects through distal regulatory elements of the genome. In this context, the three-dimensional organization of the genome is expected to play a decisive role in establishing contacts between these regulatory elements and their target genes, thereby mediating schizophrenia-associated dysregulation of gene expression. Here, we present a novel Hi-C dataset providing an unprecedented view of three-dimensional genome organization in post-mortem schizophrenia brain samples. Our findings indicate that most changes occur at long-range genomic distances while local architecture of topologically-associated domains remains largely intact. However, neurons display localized and functionally relevant loop differences, particularly in regulatory regions associated with neurodevelopmental processes. Global characteristics of higher-order chromatin organization show accelerated aging alteration pattern in schizophrenia, and downstream analysis of transcriptomic data in schizophrenia brain samples further confirms that schizophrenia is associated with accelerated aging.

## INTRODUCTION

Schizophrenia is a severe mental disorder that typically manifests in adolescence or early adulthood, often leading to functional decline and a chronic course. The incidence of schizophrenia is between 8 and 55 cases per 100,000 people per year [1], with a lifetime risk of 0.5-1% in the general population [2]. Twin studies suggest a high heritability of susceptibility to schizophrenia, estimated at 81±9% [3], although the precise genetic basis of the disorder remains largely unknown.

Many schizophrenia-associated genetic variants reside in non-coding regions and exert their effects through distal regulatory elements, such as enhancers, which interact with target genes in three-dimensional space [4–6]. During neuronal differentiation, active chromatin compartments that are maintained, as well as those newly activated, are enriched for schizophrenia risk loci [4, 7]. Recent studies have also identified neuron-specific architectural stripes, which are enriched in schizophrenia risk loci, associated with brain-specific super-enhancers, and serve as hubs for complex regulatory interactions [7, 8]. More broadly, 3D genome remodeling during the differentiation of neural progenitors affects a substantial proportion of genomic regions linked to schizophrenia susceptibility [6, 9, 10]. However, these observations were obtained based on the Hi-C maps, produced for either the healthy developing brain or neurons derived from induced pluripotent stem cells, and do not reflect chromosome conformation of the actual schizophrenia case [11]. These limitations are particularly relevant in the context of schizophrenia, where accumulating evidence suggests that chromatin architecture may interact with disease-related aging processes.

Beyond structural genome organization, schizophrenia has also been associated with broader molecular alterations consistent with disrupted biological aging. Single-cell RNA sequencing data show that aging affects all brain cell types, with interneurons being most affected, while transcriptomic effects of aging and psychiatric disorders show a strong concordance. [12–14]. These findings provide evidence for accelerated transcriptomic aging in individuals with psychiatric disorders, suggesting a potential link between transcriptomic dysregulation and the progression of age-related changes in the brain. In addition, several studies show that schizophrenia is accompanied by aging-related features at different levels of brain organization [13, 15, 16]. Further, the hypothesis of accelerated aging in schizophrenia is supported by epigenetic evidence – epigenetic clocks built on blood samples show accelerated methylome aging in schizophrenia [17, 18]. However, it remains unclear how these molecular signatures of aging are reflected in the three-dimensional genome organization of the human brain in schizophrenia.

Here, we generated a novel Hi-C dataset from postmortem brain tissue (Wernicke’s zone, Brodmann area 22) of individuals with schizophrenia (SZ) and neurotypical (HC) donors. By comparing neuronal and non-neuronal chromatin organization across diagnostic and age groups, we found that SZ is associated with altered age-dependent dynamics of long-range chromatin contacts, supporting the hypothesis of accelerated aging in SZ. To assess whether this pattern extends to another molecular layer, we constructed transcriptomic aging clocks using PsychENCODE data and observed increased transcriptomic age in individuals with SZ, further supporting the hypothesis of accelerated biological aging in this disorder.

## MATERIALS AND METHODS

### Sample collection

Post-mortem SZ and HC brain samples were obtained from the National BioService, St. Petersburg, Russia. Informed consent for the use of human brain tissues for research was obtained from all donors or their next of kin by the tissue provider bank. SZ patients were diagnosed with paranoid schizophrenia by at least two independent assessments by experienced psychiatrists. All control subjects were defined as healthy with respect to the sampled brain tissue by medical pathologists (Supplementary Table S1). All subjects suffered sudden death with no prolonged agony state from causes not related to brain function. The brain samples were collected, sliced, frozen on metal plates placed on dry ice, and stored at −80°C within 24 hours post-mortem. For sample dissection, we located the areas of interest using the Atlas of the Human Brain [19]. Prior to dissection, frozen brain slices were equilibrated to −20°C overnight. For each brain region, we dissected tissue samples of approximately 10 mg in weight using sterile and chilled using dry ice scalpels, tweezers, and tubes, and immediately stored them at −80°C. In total, 280 postmortem brain specimens were included in this study.

### Nuclei isolation

Cell nuclei were isolated from frozen brain tissue as previously described [11, 20]. Briefly, tissue (600–800 mg) was dissected, homogenized in a formaldehyde-containing buffer, fixed, and quenched with glycine. After centrifugation, nuclei were washed, further homogenized, filtered, and purified using a sucrose cushion. The nuclei pellet was resuspended, incubated with anti-NeuN fluorescent antibodies overnight, stained with DAPI, and filtered. NeuN-positive and NeuN-negative nuclei were then isolated by FANS, washed, and either stored at −80°C or used for Hi-C library preparation.

### Hi-C library preparation

The Hi-C experiment was conducted for neuronal and non-neuronal FANS-sorted nuclei extracted from six SZ and six HC samples (Supplementary Table S1), all of which were processed in parallel. However, we note that Hi-C maps derived from HC were previously published [11, 21]. Approximately 1 million nuclei per sample were extracted. The previously described protocol [21, 22] was applied in downstream steps. Hi-C libraries were sequenced on the Illumina HiSeq 4000 and Illumina Novaseq 6000.

### RNA extraction and RNA-seq library preparation

Total RNA was isolated from approximately 200 mg of brain tissue using TRIzol (Invitrogen, USA) following the manufacturer’s protocol. RNA quality was assessed with an Agilent 2100 Bioanalyzer (Agilent Technologies, USA). RNA-seq libraries were produced from 1 μg of total RNA with TruSeq Stranded mRNA kit (Illumina, USA). The libraries were sequenced on an Illumina HiSeq 4000 platform.

### Fluorescent immunostaining and imaging

Frozen brain tissue was cut in 20 μm sections on a Leica CM1950 clinical cryostat (Leica Biosystems, Germany) and thaw-mounted onto charged glass slides. The tissue sections were dried at 54°C for 30 min and further fixed in 4% paraformaldehyde for 7 min at +4°C, washed with 1x PBS, and incubated for 1 hour in blocking solutions (2.5% donkey serum, 0.5% Tween-20, 1x PBS). After blocking, the sections were incubated overnight at room temperature in the dark with Alexa Fluor 488 labeled antibodies to the neuronal nuclear marker NeuN (Abcam, UK; ab196184) in blocking solution (1:200) with 0.01% sodium azide. The stained sections were mounted in water-based Fluoromount (Sigma-Aldrich, USA) with DAPI. A LSM800 AiryScan (Zeiss, Germany) confocal microscope was utilized to image the tissue sections with Plan-Apochromat 40x/1,3 Oil DIC UV-VIS-IR objective with Zeiss Immersol 518 F oil.

### Fluorescent microscopy images processing and nuclei annotation

Fluorescence microscopy images were processed with scikit-image package v0.21.0 [23] to identify neuronal nuclei based on DAPI staining and to quantify anti-NeuN signal within segmented objects. Image stacks were converted to maximum-intensity projections, background corrected, and segmented using adaptive thresholding followed by marker-controlled watershed. Segmented regions were filtered using morphological criteria, and neuronal nuclei were identified by Otsu thresholding of mean anti-NeuN intensity (see Supplementary Methods for detailed image-processing workflow and segmentation parameters).

We analyzed 8 SZ and 8 HC cases (Supplementary Table S1). Additional samples were subsequently included in the analysis. Donors aged 41 and 45 years were classified as young. In each case 4 tissue sections were made, and 3 images were made for each section. Differences in area between groups aggregated by age and diagnosis were assessed using the Mann–Whitney U test.

### Polygenic risk scores calculation

DNA samples for genotyping were extracted from brain tissue. Genotyping was performed using the Illumina Global Screening Array (GSA) v2. Polygenic risk scores (PRS) were computed from the merged imputed genotypes (HRC r1.1, European panel) in the RMHRC dataset by downloading publicly available schizophrenia GWAS summary statistics [5], harmonising them to our SNP set and extracting the HapMap3+ variants. We computed PRS using LDpred2-auto [24], implemented in the bigsnpr package (v1.12.18) with an in-sample LD reference constructed from unrelated individuals without ancestry or heterozygosity outliers. The parameters for the “snp_ldpred2_auto” function were: shrink_corr=0.95, allow_jump_sign=TRUE, use_MLE=FALSE, burn_in=500, num_iter=500, vec_p_init=seq_log(1e-4,0.5,length.out=50); other parameters were default. All PRS were standardized to the RMHRC control group.

### Hi-C data processing

Raw sequencing reads from each Hi-C library were aligned to the human reference genome (hg38) using the distiller-nf pipeline v0.3.3 [25]. This pipeline employs BWA [26, 27] for read alignment, incorporates quality-based filtering of valid read pairs, performs genomic binning, and ultimately produces contact pair lists that are converted into binned interaction matrices. Downstream Hi-C analyses were performed on matrices merged by cell type and disease state at 5 kb resolution using cooler.merge_coolers() function of a cooler library, v0.8.11 [28]. To ensure comparability across samples, the merged contact maps were downsampled to a uniform number of contact pairs using the cooltools.sample() function from cooltools v0.5.1 [29] Final .cool matrices were normalized using ICE balancing [30] using the “cooler balance” command with its default parameters.

### General chromatin features analysis

Polymer scaling as a function of genomic distance was computed from 10-kb binned matrices using the expected_cis function from the cooltools package [29]. Scaling profiles were calculated for each sample, aggregated across autosomes, and then grouped by diagnostic cohort or donor age. For comparisons between diagnostic cohorts, mean scaling plots were calculated for each genomic distance bin and visualized on log–log axes. Cis and trans contact frequencies were computed as mean intra- and interchromosomal interaction frequencies across chromosomes and chromosome pairs, excluding chromosomes X and Y and mitochondrial sequences. The interchromosomal fraction of interactions (ICF) was defined per sample as the ratio of interchromosomal to total contacts.

### Identification and analysis of compartments

To identify compartments from Hi-C data, firstly we first applied Inspectro v0.2.0 [31] to cluster genomic bins based on the eigenvector decomposition (see Supplementary Methods for parameter settings). For each chromosome arm, we selected principal components most correlated with GC content using eigs_cis(n_eigs=5, sort_metric=‘spearmanr’) from cooltools, retaining the component best matching Inspectro clustering; in cases of ambiguity, selection was guided by visual inspection. Chromosomes Y and M and short arms (chr13p, chr14p, chr15p, chr21p, chr22p) were excluded. Differential compartment analysis between HC and SZ was performed using limma [32]. The code for saddle plots was adapted from the cooltools tutorial [33].

### Identification and analysis of TAD borders

TADs from Hi-C data were identified using the “insulation” module of the cooltools package v0.6.1 [29], which calculates the chromatin insulation profile using a diamond-window scoring approach. Insulation scores (IS) were computed at 15 kb resolution, a window size of 150 kb, a minimum valid pixel fraction of 0.75, and maintained a minimum distance of 4 bins from any bad bins. To harmonize boundary calls across samples and account for minor positional offsets, boundaries were consolidated using DBSCAN clustering algorithm from the scikit-learn package v1.5.2 [34]. Genomic coordinates of initially identified TAD borders from the Hi-C maps were organized into clusters based on proximity. Our clustering analysis utilized a resolution of 15 kb, with a clustering factor set to 2. The “cityblock” distance metric was employed, and a minimum point threshold of 3 was used to define a valid cluster. This approach allows us to identify TAD borders consistently across different samples. Boundaries co-occurring across samples within the same cluster were treated as a single consensus element in all downstream comparisons. Differences in boundary insulation strength between groups were assessed using the Mann-Whitney U test. Gene set enrichment analyses for TAD boundary-associated and non-boundary genes were carried out in R using the enrichGO function from the clusterProfiler package [35].

### Identification of chromatin loops

Chromatin loops were called using a custom analytical pipeline built around the cooler and cooltools libraries, applied to Hi-C maps aligned to the hg38 reference genome. Baseline contact frequency expectations were established per chromosome arm using the expected_cis function from cooltools, providing a null reference against which enriched interactions were evaluated. To maximize detection sensitivity across a range of interaction scales, four custom convolution kernels were designed, spanning narrow configurations for short-range contacts to broader masks targeting long-range interactions. Loop calling was performed with the dots function from cooltools, using a maximum interaction distance of 12 Mb, an FDR threshold of 0.16, tolerance for up to 5 missing bins, and a clustering radius of 2.5 bin widths. Loops identified across different kernel configurations were subsequently compared and deduplicated using custom filtering functions leveraging pairwise operations from the pybedtools package [36] retaining only high-confidence, non-redundant interactions.

### Chromatin loop processing

To reconcile minor positional discrepancies in loop coordinates across samples, loops in spatial proximity were clustered using DBSCAN (scikit-learn) and treated as consensus interactions. To reduce false negatives from cooltools loop calling, we additionally performed a supplementary search within clusters, reclassifying regions with signal intensities consistent with called loops as candidate loops and incorporating them into the final set. Resulting clusters were filtered by requiring sufficient sample support and above-threshold signal intensity. Loop anchors were intersected with genomic features in .bed/.bedpe files using pybedtools.

Differential loop analysis between SZ and HC focused on identifying up- and downregulated loops by comparing median loop intensities within clusters, with statistical significance assessed using the Mann–Whitney U test.

### Variant calling from Hi-C data and detection of variant-sensitive interactions

Variant calling from Hi-C reads aligned with distiller-nf was performed using the FreeBayes package [37]. This approach yielded 496,779 positions with variants, 468,265 of which were variable across samples. For assessment of the reliability of the variant calls, comparison with genotyping-derived variants was performed at matched positions on chromosome 18. The concordance between variants identified from genotyping and Hi-C data was 99.75%.

For detection of significant, variant-sensitive, interactions, read pairs with one read overlapping a SNP position were extracted for each sample. Only read pairs with distance of less than 1 Mb were retained for downstream analysis. Reads paired to those overlapping SNP positions were aggregated by distance from the SNP into 100-kb bins. Data from all samples were merged by variant. A generalized linear model with a Poisson distribution was then applied to obtain p-values, accounting for GC content, mappability, fragment size, expression in the interacting bin, and the difference in PC1 values between the interacting regions. An interaction was defined as allele-specific if it was significant for one variant (adjusted p-value < 0.05) but not for another (adjusted p-value > 0.1).

The annotation of interacting bins or SNP positions was based on the intersection with coordinates of genes (full length). Primary eQTL–gene linkages were retrieved from the MetaBrain project (european population, cortical data) [38].

Details of the analysis, additional filtering, and instrument parameters applied are described in the Supplementary Materials.

### Analysis of heterozygous inversion

The inversion was identified by visual inspection, with breakpoints defined at 10 kb resolution. The control sample (HC-2) was selected based on a comparable total number of contacts in the initial neuronal Hi-C maps (104.6 million in HC-2 versus 106.8 million in SZ-1). The intensity of the loop extrusion track in both control and affected samples was calculated from fragments of the Hi-C map corresponding to this chromatin feature, and then normalized by the average contact frequency within a stripe located downstream of the breakpoint (Supplementary Figure S1).

### Transcriptomic data processing and differential gene expression analysis

Trimmed FASTQ files were processed using Salmon v1.10.3 [39]. GENCODE v46 gene annotations were used to obtain per-gene counts with the tximport package [40]. DESeq2 [41] was applied to generate rlog-transformed values from raw counts. To account for differences in cell composition that could confound gene expression comparisons, we performed cell-type deconvolution on rlog-transformed values, using the CIBERSORT package [42], with gene signatures based on Human Cell Atlas transcriptomic data sourced from Sutten et al. [43].

Publicly available RNA-seq data on sorted human brain nuclei (NeuN-positive and NeuN-negative) [44] were processed as follows: FASTQ files were analyzed using the nf-core/rnaseq pipeline v3.8.1 [45, 46] with parameters --aligner hisat2, --genome hg38. Resulting BAM files were sorted by name using samtools sort (v1.6) [47]. The transcriptional profile for the heterozygous inversion analysis was generated using the bamCoverage tool from the deepTools package [48]. Abundance used in the analysis of variant-sensitive contacts was obtained with the Salmon and tximport packages.

### Transcriptomic aging clocks construction and differential gene expression analysis of data from PsychENCODE consortium

Publicly available transcriptomic data for post-mortem bulk brain tissue from PsychENCODE consortium [49, 50] were used to estimate the contribution of SZ to the aging process. The raw read counts were normalized with pyDESeq2 package v0.4.4 [51]. Genes with at least ten read counts on average were kept. Two variables, age group and diagnosis, were added to pyDESeq2 design formula. Differentially expressed genes with adjusted p-value < 0.1 (Benjamini-Hochberg correction) were used to assess the codirectionality of gene log2FoldChange between age and diagnosis conditions.

Transcriptomic aging clocks were constructed using Elastic Net (scikit-learn package) regression to predict age from gene expression data. Model performance was evaluated using fivefold cross-validation, with regularization parameters set to α = 1.27 and l1_ratio = 0.9. Age acceleration was defined as the difference between predicted and chronological age for each individual. The Mann-Whitney U test was conducted to validate the statistical significance of age acceleration between the two health conditions.

### Age Score construction

Publicly available RNA-seq data on sorted human brain nuclei [52, 53] were used to construct and evaluate Age Score metrics across diagnoses. Gene expression data were normalized with pyDESeq2 using variance stabilizing transformation (VST).

To construct the Age Score, expression values of the selected genes were first scaled to reduce the influence of extreme values. For each gene *g* and sample *i*, normalized expression 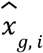 was computed by dividing raw VST expression *x*_*g, i*_ by the gene-specific 95th percentile, followed by scaling with the median absolute deviation.

The final Age Score for the sample *i* was defined as:

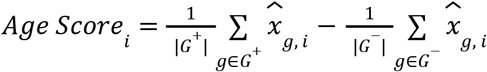

where *G*^+^ and *G*^−^ denote the sets of genes with positive and negative age-associated regression coefficients, respectively. An ordinary linear model was introduced to analyze Age Score distribution, the next model design was used: Age Score ∼ Age * Diagnosis.

### Chip-seq data processing and analysis

To annotate the heterozygous inversion, ChIP-seq data for CTCF in neurons and oligodendrocytes from dorsolateral prefrontal cortex were obtained from the BrainTF resource [54]. Corresponding bigWig tracks were generated using the nf-core/chipseq pipeline [55] with the hg38 genome assembly. BigWig files were converted to bedGraph format, and average signal intensity was computed for each 10-kb genomic bin. Replicates were merged by calculating the mean signal and input in each bin. Next, data were normalized by dividing the signal track by its input track, with a pseudocount defined as the minimum non-zero value.

## RESULTS

### Hi-C profiling of schizophrenia and control brain samples

To investigate the fine-scale details of chromatin structure in SZ-affected brain tissue, we collected superior temporal gyrus samples (Wernicke’s area, Brodmann area 22) from six SZ cases with age distribution and sex composition similar to six HC cases (Figure 1A) [11, 21]. As group assignment was based solely on clinical psychiatric diagnosis rather than laboratory measures, we additionally obtained genotyping data for 10 of 12 samples and calculated polygenic risk scores (PRS; Figure 1B). Although individual samples show some divergence between diagnosis and PRS, SZ samples tend to have higher PRS values than HC, confirming the validity of our diagnostic grouping. Assuming that neurons and glia might contribute to SZ through parallel yet distinct mechanisms, we isolated these populations via fluorescence-activated nuclei sorting (Figure 1C). Separately for neuronal and non-neuronal nuclei, we generated Hi-C maps with contact numbers ranging from 78 to 156 million (Figure 1D); after downsampling, this enabled analysis at up to 15-kb resolution.

**Figure 1.**
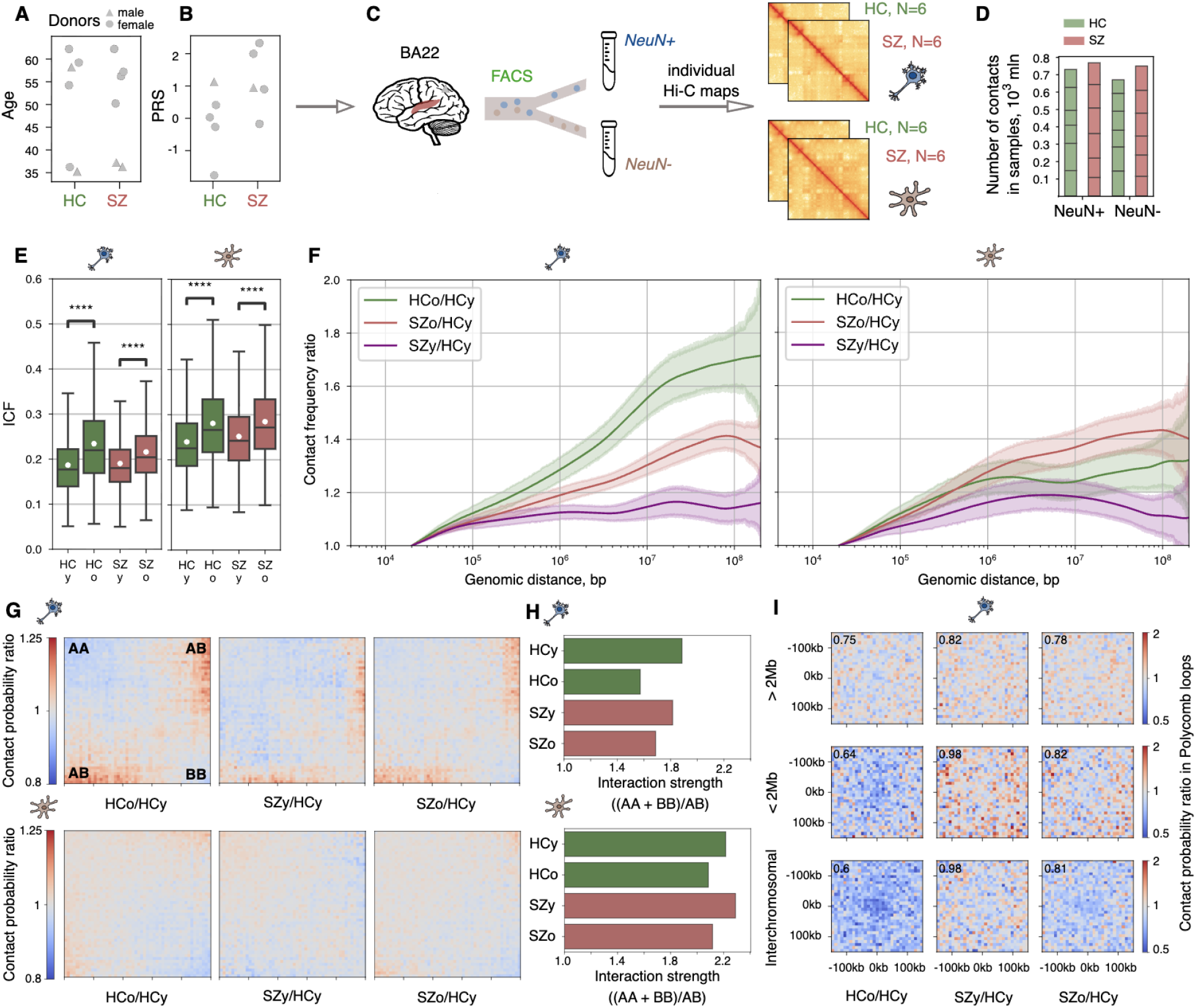
Large-scale chromatin features in SZ and HC. **A-B**. Characteristics of diagnostic cohorts: age and sex (A), polygenic risk score (B). **C**. Anatomical localization of the analyzed brain region, experimental procedure and the design of this study. **D**. The number of contacts in samples. **E**. ICF for young (y) and old (o) donors calculated for every 50-kb genomic region for HC (green) and SZ (red) in neurons (left) and non-neurons (right). White dots represent mean values. Asterisks indicate Mann-Whitney U test p-value: **** - p < 0.0001. **F**. Average polymer scaling ratios calculated for HCo/HCy (green), SZy/HCy (purple) and SZo/HCy (red) in neurons (left) and non-neurons (right). **G**. Average saddle plot ratios calculated for HCo/HCy (left), SZy/HCy (middle) and SZo/HCy (right) in neurons (top) and non-neurons (bottom). **H**. Compartment interaction strength (AA+BB)/AB calculated within top-20% areas of saddle plots in neurons (top) and non-neurons (bottom). **I**. HCo/HCy (left), SZy/HCy (middle) and SZo/HCy (right) average observed/expected Hi-C signal of Polycomb loops in neurons. The number in the corner corresponds to the central pixel.

### Large-scale chromatin features show accelerated aging in schizophrenia

Having established the overall comparability of the obtained Hi-C maps (Supplementary Figure S2A-D), we first asked whether SZ is associated with global changes in higher-order chromatin organization. Given previous evidence for accelerated aging in SZ relative to HC [13, 14], we specifically examined whether age-associated chromatin changes differ between diagnostic cohorts. We stratified samples into young (< 40 years; 2 HC and 2 SZ samples, further referred to as HCy and SZy) and old (≥ 50 years; 4 HC and 4 SZ samples, further referred to as HCo and SZo) groups, and compared age-dependent changes in global chromatin features between HC and SZ.

In both neuronal and non-neuronal nuclei, analysis of the interchromosomal fraction of interactions (ICF) revealed a significantly higher proportion of interchromosomal contacts in the old group than in the young in both diagnostic cohorts (Figure 1E, Mann-Whitney U test, p-value < 1×10^-4^). Notably, SZy showed a shift toward higher ICF values compared with HCy in both nuclei populations, placing SZy closer to the range observed in older donors. This observation suggests that disease-associated changes in higher-order chromatin organization are already detectable in young SZ samples and partially resemble age-associated changes observed in HC.

To determine whether this pattern extends to distance-dependent intrachromosomal contacts, we next analyzed age- and diagnosis-related changes in polymer scaling profiles. Using HCy as a baseline, we calculated average HCo/HCy, SZy/HCy, and SZo/HCy ratios of polymer scaling. In both neuronal and non-neuronal nuclei, SZy shifted toward the profiles of older donors in the range of long-distance contacts (Figure 1F), suggesting that young SZ chromatin already adopts a more aged-like configuration.

We next examined whether these aged-like shifts in global contact patterns were accompanied by changes in A/B compartment organization. Sample-to-sample correlations of compartment eigenvectors showed that SZ samples were not more heterogeneous than HC samples (Supplementary Figure S2E, F). We identified eight genomic regions with significant differences between HC and SZ, but only one showed a noticeable effect size upon visual inspection. This region contains the *CELF* gene, where eigenvector values were higher in HC than in SZ, while contact frequency was elevated in SZ (Mann-Whitney U test, p-value = 7.38×10^-5^), indicating a more compacted heterochromatin state (Supplementary Figure S2G-I).

Next, we visualized age- and diagnosis-related changes in compartment strength using HCo/HCy, SZy/HCy, and SZo/HCy ratios of average observed/expected Hi-C contact probability as saddle plots, in which genomic regions are arranged by the corresponding PC1 rank (Figure 1G). In both neuronal and non-neuronal nuclei, interactions between compartments (AB) increased in older donors in both diagnostic cohorts, accompanied by a parallel decrease in within-compartment interactions (AA and BB). These coordinated changes were captured by compartment interaction strength, calculated as (AA+ BB)/AB (Figure 1H), and indicate weaker compartmentalization in older donors of both diagnostic groups. In neurons, the strongest reduction in compartmentalization was observed between HCy and HCo, whereas SZy showed an intermediate compartment interaction strength between HCy and HCo, again consistent with an aging-like shift in young SZ samples.

Finally, we focused on neuronal Polycomb loops, a class of neuron-specific long-range contacts enriched for H3K27me3 at their anchors, which we described in our previous work [21]. The anchors of Polycomb loops are strongly enriched for developmental transcription factors, suggesting potential relevance to SZ pathogenesis. Using previously published Polycomb loops annotations [56], we quantified average loop intensities and assessed their age- and diagnosis-related changes, again using HCy as a baseline (Figure 1I). In HCo neurons, Polycomb loop intensity declined substantially relative to HCy, while SZo showed a smaller decline and SZy showed only a slight decrease relative to HCy. Nevertheless, the direction of change in SZy was shifted toward SZo, consistent with accelerated aging-like chromatin changes in SZ.

### Fine-scale chromatin architecture is largely preserved in schizophrenia

We next investigated whether these accelerated aging-like patterns of chromatin changes extend to the level of topologically associating domains (TADs). TAD numbers (Supplementary Figure S3A, B), boundary position similarity (Supplementary Figure S3C, D), and average interaction intensity within TADs and at TAD boundaries (Supplementary Figures S3E, F, S4, S5) were broadly comparable between SZ and HC in both neuronal and non-neuronal nuclei. Consistently, PCA of insulation scores showed no clear separation between diagnostic groups in either nuclei population (Supplementary Figure S6A, B). Although differential boundary analysis identified altered boundaries, genes located within these regions showed no functional enrichment, suggesting that these differences lack functional coherence. We also found no robust evidence that TAD-level alterations in SZ resemble age-related alterations in HC (Fisher’s exact test for overlap between SZ-associated and age-associated differential boundaries, p-value > 0.05). Together, these analyses indicate that TAD organization is largely preserved in SZ. Importantly, HC and SZ showed comparable within-group heterogeneity (Supplementary Figure S3C,D), suggesting that the lack of SZ–HC separation is not driven by greater variability in one cohort.

We next asked whether we could trace any accelerated aging-like patterns at the level of chromatin loops. Similar to TADs, the total number and length distribution of loops (Supplementary Figure S3G, H), loop positions (Supplementary Figure S3I, J) and their average intensity (Supplementary Figure S3K,L) did not differ significantly between SZ and HC in either neuronal or non-neuronal nuclei. Variability within HC and SZ was comparable (Supplementary Figure S3I,J), suggesting that this lack of differences is not driven by greater variability in one cohort. Yet, although PCA of loop intensities across individual samples revealed no separation between SZ and HC in non-neuronal nuclei (Supplementary Figure S7), neuronal samples separated by diagnosis along PC2, suggesting a degree of neuron-specific loop-level distinction worth further exploration (Figure 2A). Consistently, cluster analysis of loop intensities across HCy, HCo, SZy, and SZo groups showed that, only in neurons, older samples (HCo and SZo) had the most similar loop intensity profiles, while SZy occupied an intermediate position between HCy and older samples of both diagnostic groups (Figure 2B). This pattern is consistent with the accelerated aging-like changes observed in SZy at higher-order levels of chromatin organization.

**Figure 2.**
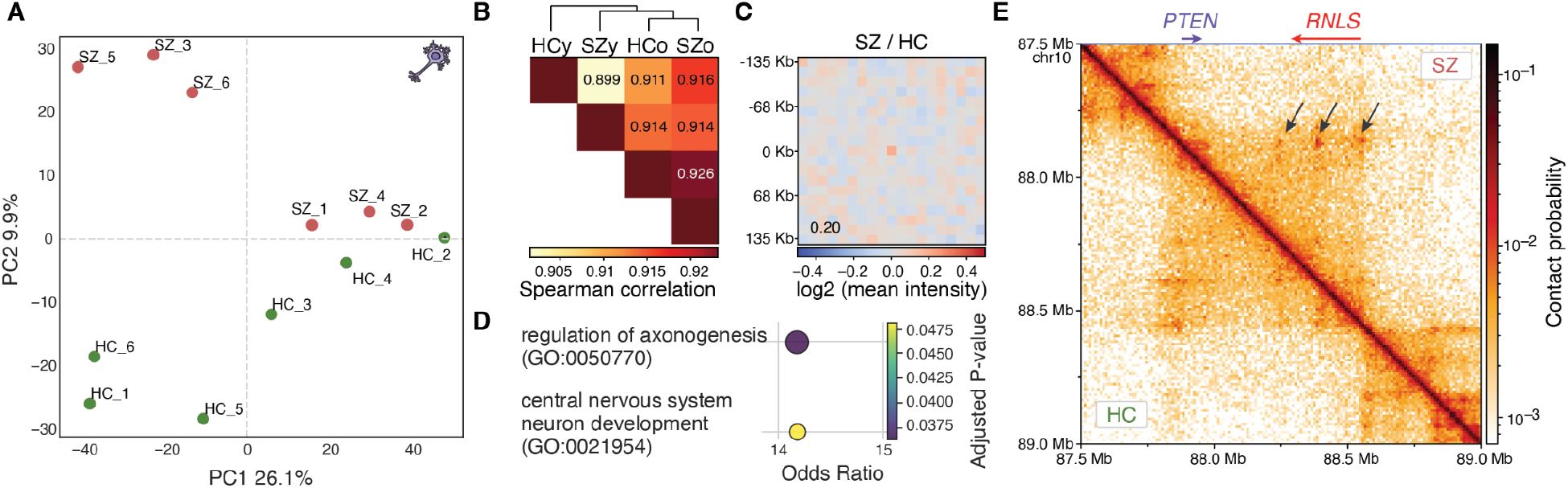
Analysis of TADs and chromatin loops in neuronal cells. **A**. PCA of loop intensities across analyzed samples for neurons. **B**. Hierarchical clustering and Spearman correlations of chromatin loop intensities across four sample groups separated by diagnosis and age (HCy, HCo, SZy, SZo). **C**. Average loop intensity ratio (SZ to HC) for loops within the top 10 quantile by feature importance (PC2 of loop intensity, A). **D**. GO enrichment and loop intensity ratio analyses for genes whose promoters overlap neuronal loop anchors within the top 10 quantile by feature importance. **E**. Differential chromatin loops at the *PTEN* gene promoter in neurons.

To identify specific chromatin loops driving the separation along the PC2 in neurons, we ranked loops by their importance to PC2 separation and focused on the top 200 most influential loops. Annotation of gene promoters at these loop anchors followed by GO enrichment analysis revealed that loops elevated in SZ neurons were significantly enriched for the “regulation of axogenesis” GO term (Figure 2C, D). Several genes within this term, including *PTEN*, not only have documented associations with SZ but also show visually discernible differential loop interactions in our Hi-C data, exemplified by two distinct loops involving the *PTEN* locus (Figure 2E). Overall, these results indicate that neurons display localized and functionally relevant loop differences, particularly in regulatory regions associated with neurodevelopmental processes, warranting further study into their potential role in the pathophysiology of SZ.

### Fluorescent microscopy revealed preservation of nuclear size in schizophrenia

Because local structures – TADs and most chromatin loops – were largely preserved in SZ, whereas long-range chromatin interactions showed disease- and age-associated alterations, we asked whether these changes could reflect differences in nuclear morphology. Specifically, we hypothesized that altered nuclear size might contribute to the observed changes in higher-order chromatin organization in SZ. To test this, we performed fluorescence microscopy of neuronal nuclei stained with DAPI and anti-NeuN antibodies (Figure 3A-C).

**Figure 3.**
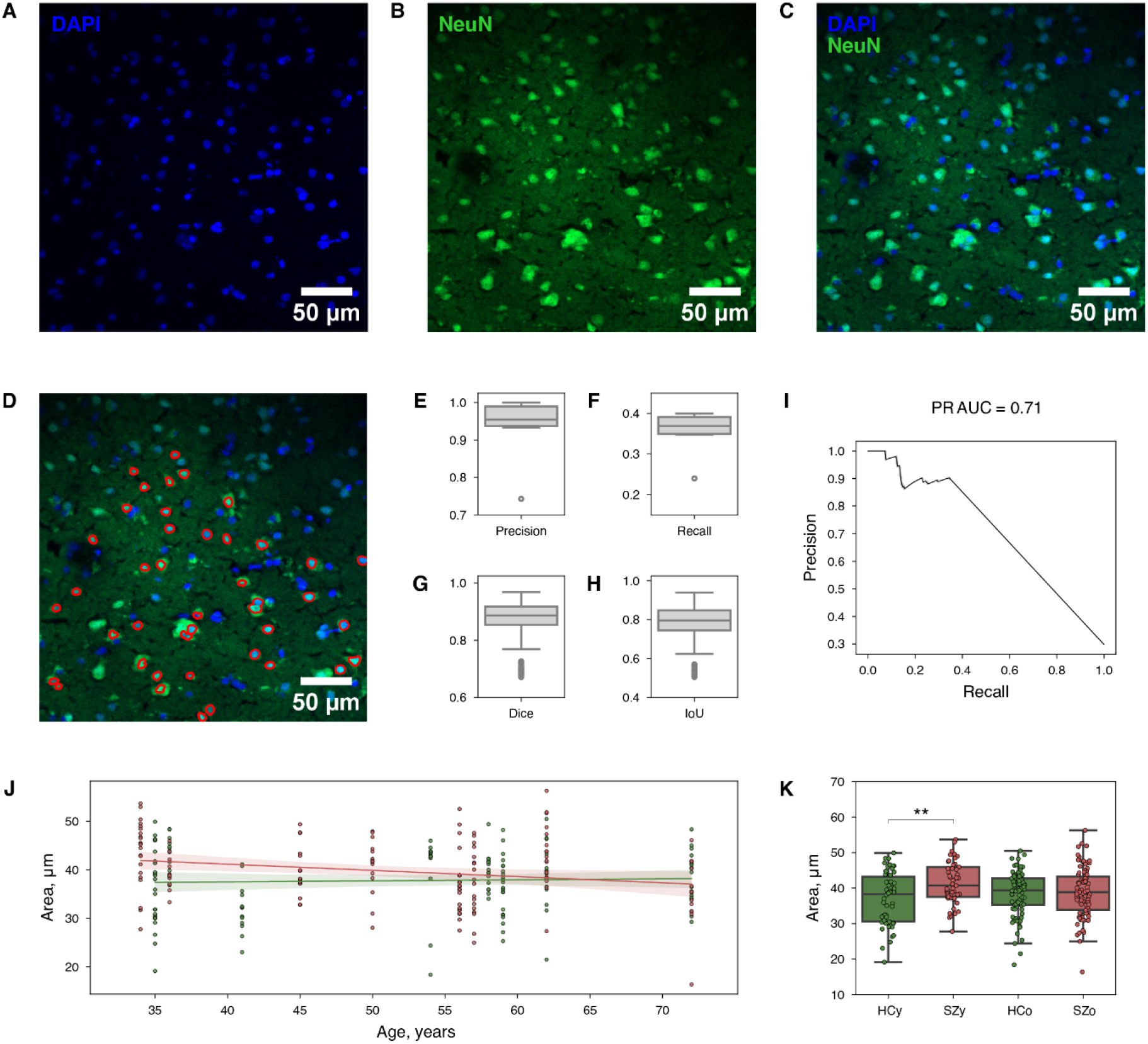
Analysis of brain tissue microscopy and nuclei sizes. **A-C**. Nuclei staining for double-stranded DNA with DAPI and with antibodies against the neuronal marker NeuN. **D**. A semi-automated segmentation pipeline was applied to select only neuronal nuclei (red contour). **E-H**. Accuracy of the pipeline performance: precision (E), recall (F), Dice (G), and IoU segmentation (H) metrics. **I**. PR curve for the pipeline performance. **J**. The age-related dynamics of the nuclear area in HC (green) and SZ (red). **K**. Comparison of nuclear area between age groups and diagnostic cohorts. Asterisk indicates Mann-Whitney U test p-value: ** - p-value < 0.01.

Neuronal nuclei were identified from fluorescence microscopy images using a semi-automated segmentation pipeline (Figure 3D). The pipeline achieved an average precision of 0.93 (Figure 3E). Recall was more moderate, reaching 0.35 (Figure 3F), primarily because large nuclear aggregates could not be reliably separated and were therefore excluded. We considered this recall acceptable because this approach guaranteed segmentation purity. Segmentation quality was further evaluated using the Dice similarity coefficient and Intersection over Union (IoU), which quantify the overlap between predicted and manually annotated nuclear masks (Figure 3G, H). This method achieved a Dice score of 0.87 and an IoU of 0.77, indicating strong agreement with manual segmentation. Additionally, the precision–recall area under the curve (PR AUC) reached 0.71 (Figure 3I), supporting acceptable performance under class-imbalanced conditions.

We next analyzed whether the nuclear size differed between SZ and HC. Quantitative analysis of the neuronal nuclear area revealed a significant increase in SZ samples compared with HC (Figure 3J, K). The mean nuclear area was 41.40 μm^2^ in SZ versus 36.94 μm^2^ in HC, corresponding to an approximately 12% increase (Mann–Whitney U test, p-value = 3.9×10^-3^). These results suggest that altered nuclear size may contribute to the observed changes in long-range chromatin interactions in SZ.

### Gene expression analysis evidenced accelerated aging in schizophrenia

To validate and estimate age acceleration in SZ, we reanalyzed publicly available transcriptional data from the PsychENCODE project. Transcriptional age was predicted by training an Elastic Net regression model with fivefold cross-validation. Comparison of the chronological and transcriptional ages in the test dataset revealed an increase of 8.03 years (Mann-Whitney U test, p = 3.4×10^-5^) of age acceleration rate in SZ compared to HC (Figure 4A, B).

**Figure 4.**
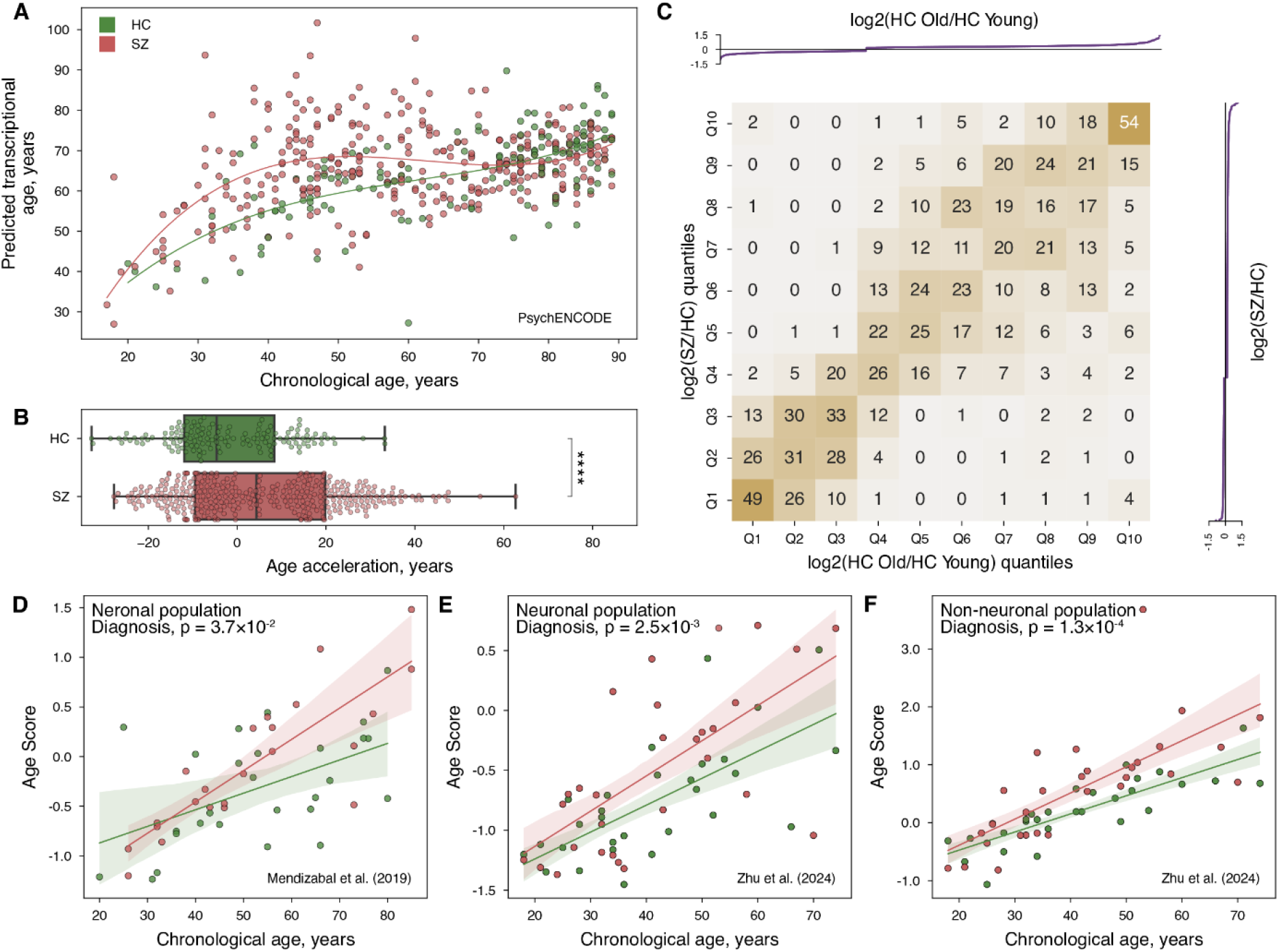
Analysis of transcriptional age acceleration. **A**. Elastic Net-based prediction of transcriptional age over chronological age. **B**. Aging acceleration in bulk tissue according to PsychENCODE data. Asterisks indicate Mann-Whitney U test p-value: **** - p-value < 0.0001. **C**. The heatmap, indicating a strong concordance in the directionality of expression changes between the two factors of age and diagnosis. **D-F**. The Age Score was constructed based on three publicly available datasets in both neuronal and non-neuronal cells. The fixed effect of diagnosis was calculated using ANOVA, Diagnosis factor.

To investigate the relationship between gene expression changes associated with age and disease status, a differential expression analysis was performed. Significantly altered genes for each factor were independently identified, and their expression changes were stratified into deciles according to the log2FC values (Figure 4C). The resulting gene distribution over deciles revealed a strong concordance in the directionality of expression changes between the two factors. Specifically, genes that were upregulated in older donors tended to be similarly upregulated in disease, while downregulated genes exhibited a corresponding analogous pattern. This observation suggests a substantial overlap between age-related and disease-related transcriptional responses.

Additionally, we performed a reanalysis of publicly available datasets for sorted neuronal and non-neuronal cells [52, 53]. To assess the contribution of SZ to aging, we scored age-related transcriptional changes for SZ and HC (see Methods). The Age Score reflects the balance between gene expression programs that increase and decrease with age: higher values indicate a shift toward an “older” transcriptional profile, while lower values correspond to a more “youthful” state. Intuitively, it summarizes how strongly a sample resembles the gene expression pattern typically associated with aging. A linear model was used to evaluate the relationship between the obtained Age Score and chronological age, diagnosis, and their interaction. In all three cases, there was a significant effect of diagnosis (p-value < 0.05), indicating higher parameter values in individuals with SZ relative to HC (Figure 4D-F).

Overall, reanalysis of transcriptomic data revealed a significant increase in aging rates in individuals with SZ. Gene expression changes associated with aging showed strong concordance with disease-related transcriptional alterations, suggesting overlapping molecular programs between aging and SZ.

### Variant-sensitive chromatin interactions

As Hi-C data are generated by sequencing DNA fragments and therefore contain genomic information, we used these data to investigate allele-specific contacts in neurons. We performed variant calling on the Hi-C data and identified sites covered in any sample by at least 30 reads (see Methods). We then detected significant allele-specific contacts between these SNP positions and 100-kb bins located no more than 1 Mb apart (Figure 5A, see Methods). The large bin size was chosen because of the sparsity of pairwise contacts. Significance of interactions was assessed using a generalized linear model with Poisson distribution. In addition to GC content and fragment size, we accounted for gene expression levels and differences in compartment activity between interacting loci. Most comparisons were conducted between heterozygous and homozygous variants, as the majority of genetic variants across samples was presented by such a combination. This approach yielded 103 significant, allele-specific, contacts (Figure 5B, Supplementary Table S2), most of which occurred between regions containing two different genes, within the same gene, or loci with SNPs located in intergenic regions and genes (Figure 5C). As confirmation of the validity of these contacts, we found among them an interaction between rs4835690 and a bin containing the *MATR3* gene, located at a distance of nearly 700 kb from the SNP. This gene was previously reported to be upregulated upon mutagenesis of a repressive sequence located at approximately the same distance (695 kb), with which it interacts [10].

**Figure 5.**
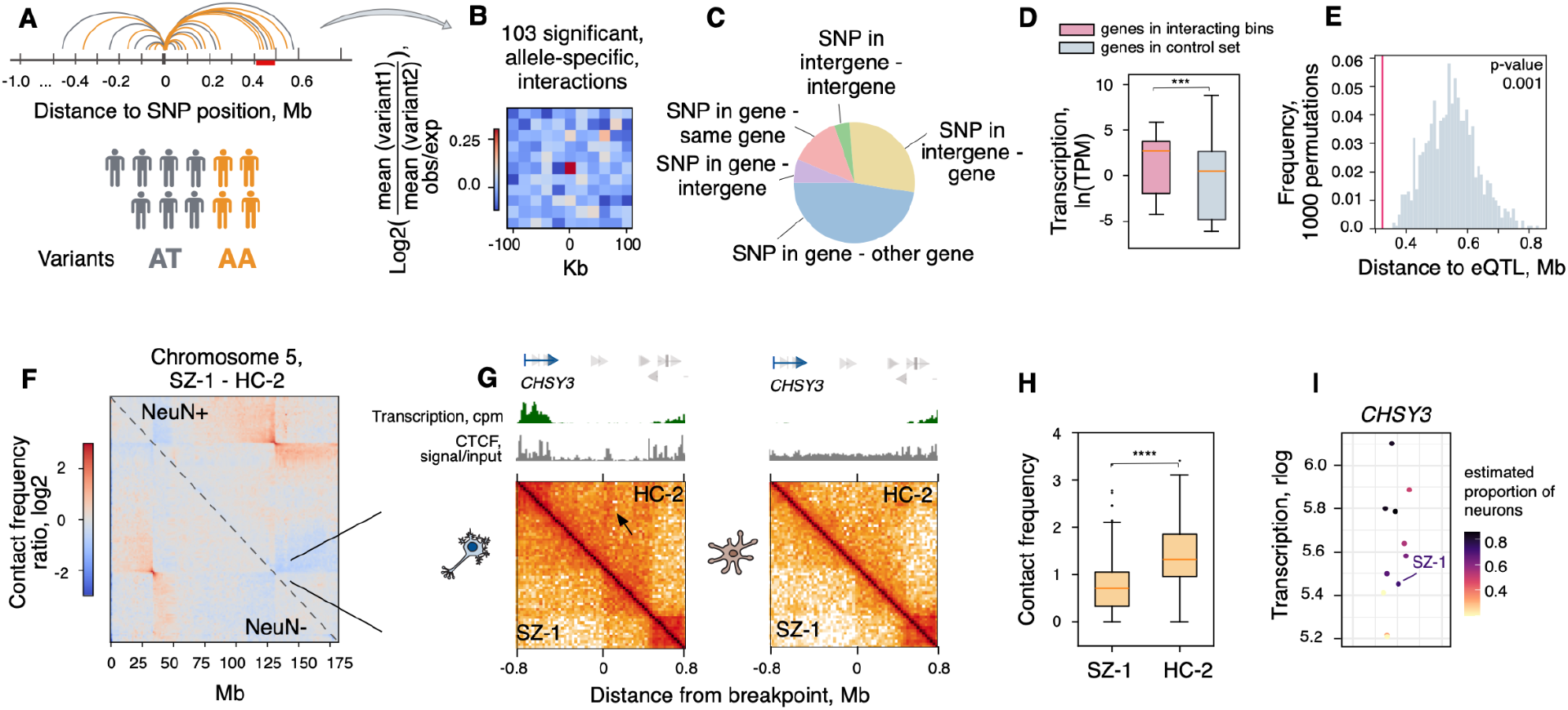
Variant-sensitive chromatin interactions. Heterozygous inversion. **A**. Design of the approach to detect significant allele-specific contacts. **B**. Differences in allele-specific interactions, averaged across samples with the same variant. **C**. Annotation of detected contacts. **D**. Transcription of genes intersecting 100 kb bins which interact with SNP locus in allele-specific manner. Pink: genes in significant, variant-sensitive, interactions; light blue - control set of genes. **E**. Distance distribution from primary eQTL of genes to interacting SNPs in allele-specific contacts (light blue histogram: distance-aware control; pink: observed contacts). **F**. Hi-C maps of chromosome 5 in SZ-1 sample with heterozygous inversion (upper right: neurons; bottom left: glia). **G**. Hi-C maps, transcription profiles, and CTCF ChIP-seq tracks around the right breakpoint in neurons (left) and non-neuronal cells (right). The black arrow points to the extrusion track in the unaffected sample. **H**. Contact frequency in area corresponding to extrusion track. I. *CHSY3* gene expression across samples. Asterisks indicate Mann-Whitney U test p-values: *** - p-value < 0.001 (E) and **** - p-value < 0.0001 (H).

To annotate genes involved in variant-sensitive contacts with SNPs, we leveraged public data on sorted neuronal transcriptome [44] and revealed that the genes involved in interactions with SNPs showed significantly higher expression than the control gene set (Mann-Whitney U test, p-value = 1.2×10^-4^, Figure 5D, see Methods). We also used data on linkage between primary eQTLs and their respective genes from the MetaBrain project [38] and demonstrated that the distance between SNP positions and eQTLs of genes involved in these variant-sensitive contacts was significantly shorter than in a random, distance aware, control (permutation p-value 0.001, Figure 5E). Among self-interacting genes, most of which were, as expected, among the top decile in gene length (mean length approximately 720 kb), we identified several genes associated with psychiatric disorders according to GWAS analysis, including *ZNF804A, DPYD, CACNA1C*, and *SATB2*. Because the exact mechanisms by which these genes and genomic variants contribute to neuropathogenesis remain unclear, contacts within these genes that may have a regulatory effect merit further attention and investigation.

Notably, this analysis does not reveal the true causality or regulatory impact of the variants involved in the detected contacts. Due to the uneven coverage inherent in Hi-C data, not all genomic variants can be reliably detected or analyzed; therefore, causal variants might be missed, despite being in high linkage disequilibrium with those examined. Nevertheless, the detected SNP pairs, as well as variants in high linkage disequilibrium with them, and the genes involved in allele-specific contacts may be useful in future studies of how variants affect gene regulation through spatial chromatin organization.

Hi-C data also provide information on large-scale structural variants. Visual inspection of individual Hi-C maps revealed a characteristic “butterfly” pattern [57], corresponding to a 96.5 Mb heterozygous inversion on chromosome 5 in one SZ sample (Figure 5E). One of the inversion breakpoints mapped to a repetitive region (34.19 - 34.2 Mb) containing the *C1FQTNF3* gene, where low coverage complicated further analysis. The other breakpoint is located in an intergenic region (130.68 - 130.69 Mb) and falls within the inner part of a TAD. This breakpoint seemingly disrupts interactions between CTCF-associated regions that form a loop in the control NeuN-positive sample, leading to a decrease in the prominence of the extrusion track (Mann-Whitney U test, p-value = 5.1×10^-22^, Figure 5H, Supplementary Figure S1). One loop anchor contains the TSS of the *CHSY3* gene, which shows moderate expression in neurons and low expression in glial cells. Analysis of our sample-matched bulk transcriptome, after accounting for estimated cell composition, showed the lowest expression in the sample carrying the inversion among all samples with a similar estimated proportion of neurons (Figure 5I).

Taken together, this finding suggests a possible indirect effect of structural variation on gene expression at the regulatory level via disruption of chromatin architecture, which may additionally contribute to genetic predisposition to psychiatric disorder.

## DISCUSSION

Despite efforts to characterize the regulatory genome in SZ, a fundamental limitation that persisted across prior studies is reliance on interaction data derived exclusively from neurotypical individuals [6, 7, 10]. This usage of neurotypical 3D genome architectures assumes that chromatin organization is largely preserved in SZ, leaving the structural genomic landscape of the disorder itself uncharacterized. Here, we address this gap by generating the first Hi-C maps directly from postmortem brain tissue of individuals diagnosed with SZ for both neuronal and non-neuronal cell types. This dataset provides an unprecedented view of three-dimensional genome organization in the SZ brain, revealing widespread alterations in higher-order chromatin structure that are invisible to analyses anchored in neurotypical reference data.

Our findings indicate that SZ is associated with an early shift toward an aged-like chromatin state. At the fine scale, changes were limited: TAD organization was largely preserved, and only neuronal loop intensities showed patterns consistent with accelerated aging. At the large scale, chromatin from SZy samples showed aged-like patterns in ICF, polymer scaling, compartment interaction strength, and Polycomb-mediated loop intensity. Together, these results suggest that SZ is associated primarily with early aging-like alterations in large-scale chromatin organization, while fine-scale chromatin architecture remains comparatively preserved.

This architectural evidence for accelerated aging in brain cells is supported by our transcriptomic analyses, which identified a transcriptional age acceleration of approximately 8.03 years in SZ. Previous studies have shown that SZ is accompanied by molecular signatures associated with accelerated aging, including strong concordance between age-related and disease-associated transcriptomic changes in the brain, as well as accelerated methylome aging detected by epigenetic clocks [12–14, 17, 18, 58]. Using an Age Score approach in sorted cell populations, we further found evidence of accelerated aging-related transcriptional changes in both neuronal and non-neuronal cells. Consistently, our Hi-C analyses indicate that large-scale chromatin features show aging-like shifts in both nuclear populations. At the fine scale, however, TAD organization was largely preserved in both neuronal and non-neuronal nuclei, whereas loop-level alterations were only detectable in neurons; these changes were localized and subtle, yet involved genes associated with regulation of axogenesis, including the risk genes.

Because nuclear geometry can influence long-range interaction frequencies, we asked whether altered nuclear size might contribute to the higher-order chromatin changes observed in SZ. SZ samples showed a statistically significant 12% increase in neuronal nuclear area compared with HC. These results suggest that altered nuclear size likely contributes to the observed changes in long-range chromatin interactions in SZ, although additional molecular mechanisms affecting higher-order chromatin organization may also be involved.

Our study also leveraged Hi-C data to explore the intersection of genetic variation and 3D structure. We identified over 100 significant allele-specific interactions in neurons. The proximity of these interacting SNPs to known expression quantitative trait loci supports the functional relevance of these contacts. A compelling example was the interaction between the *MATR3* gene and a distal SNP, which mirrors previously reported repressive regulatory mechanisms [10]. Furthermore, the identification of a 96.5 Mb heterozygous inversion on chromosome 5 in one SZ sample allowed us to observe the direct impact of structural variants on chromatin extrusion. The disruption of a CTCF-associated loop at the inversion breakpoint was associated with the downregulation of the *CHSY3* gene, providing a clear example of how large-scale genomic rearrangements can rewire the regulatory landscape.

By providing the first Hi-C maps generated from SZ brain tissue, this study demonstrates that the disorder is characterized not by a global breakdown of chromatin domains, but by a subtle rewiring of neurodevelopmental loops and an acceleration of the aging process at the level of large-scale chromatin features. These results shift the focus toward the temporal dynamics of genome organization, opening new avenues to investigate whether modulating these accelerated aging signatures could slow the progression of the disease.

## Supporting information

Supplementary Table S1. Sample metadata

Supplementary Table S2. Significant variant-sensitive interactions

## DATA AVAILABILITY

Sequencing data generated in this study are available in the NCBI GEO database under the accession numbers GSE330387 and GSE330768. Other datasets analyzed in this study were obtained from publicly available domain resources. The Hi-C data on sorted human brain nuclei of neurotypical donors are available in NCBI GEO under accession numbers GSE229816 and GSE291967. The RNA-seq data on sorted human brain nuclei are available in NCBI GEO under accession numbers GSE174407, GSE107638, and GSE96615. The RNA-seq data on bulk brain tissue and ChIP-seq data for CTCF in neurons and oligodendrocytes were obtained from the PsychENCODE Consortium at https://doi.org/10.15154/1g4m-dy13. Genetic data are available through the “Neuroresource” collection (https://ckp-rf.ru/catalog/usu/4145280/) of RMHRC. Microscopy images generated and analyzed in this study are publicly available through the Zenodo dataset at https://doi.org/10.5281/zenodo.20098460.

## ACKNOWLEDGEMENTS

We are grateful to the Skoltech BioImaging and Spectroscopy Core Facility for the excellent technical assistance. We thank the Center for Precision Genome Editing and Genetic Technologies for Biomedicine, IGB RAS for the cell sorting. We thank Dr. Dmitrii Kriukov and Evgeniy Efimov for helpful advice, Skoltech Genomics Core Facility and Dr. Elena Shagimardanova (Skolkovo Institute of Science and Technology) for sequencing and quality control of Hi-C libraries, and the Shared Research Facility “Shared Research Center of the Ivannikov Institute for System Programming of the Russian Academy of Sciences (SRC ISP RAS)” for providing computational equipment.

## Author contributions

Kirill A. Ulianov: Investigation, Formal analysis, Writing—original draft, Writing—review & editing. Diana R. Zagirova: Formal analysis, Writing—original draft, Writing—review & editing. Anna D. Kononkova: Formal analysis, Writing—review & editing. Anastasiia Dudkovskaia: Formal analysis, Writing—original draft, Writing—review & editing. Maria N. Molodova: Investigation, Writing—review & editing. Kirill V. Morozov: Formal analysis, Writing—review & editing. Olga I. Efimova: Investigation, Formal analysis, Writing—review & editing. Maria Bazarevich: Investigation, Formal analysis, Writing—review & editing. Alexander V. Cherkasov: Investigation, Formal analysis, Writing—review & editing. Polina D. Morozova: Investigation, Writing—review & editing. Anna V. Tvorogova: Investigation, Writing—review & editing. Ilya A. Pletenev: Formal analysis, Writing—original draft, Writing—review & editing. Nikolay V. Kondratyev: Resources, Formal analysis, Writing—review & editing. Vera E. Golimbet: Resources, Formal analysis, Writing—review & editing. Sergey V. Razin: Investigation, Writing—review & editing. Philipp Khaitovich: Resources, Writing—review & editing. Sergey V. Ulianov: Investigation, Supervision, Writing—original draft, Writing—review & editing. Ekaterina E. Khrameeva: Conceptualization, Formal analysis, Supervision, Writing—original draft, Writing—review & editing.

