## Supplementary Materials for "Accelerated Aging Signatures in 3D Genome Organization and Transcriptome in Schizophrenia"

### SUPPLEMENTARY METHODS

##### Fluorescent microscopy images processing and nuclei annotation

Fluorescence microscopy images were processed with scikit-image package v0.21.0 [[1]](https://www.zotero.org/google-docs/?9Vja0A) to identify neuronal nuclei based on DAPI staining and to quantify anti-NeuN signal within segmented objects. Raw image stacks were loaded and reduced to two-dimensional images using maximum intensity projection for each channel. All images were converted to grayscale. To correct for uneven illumination, background subtraction was applied using a Gaussian filter, followed by intensity rescaling. Nuclear candidates were then segmented using adaptive thresholding (Sauvola method), which enables robust binarization under spatially varying background conditions.

To separate touching nuclei, a distance transform was computed from the binary mask and smoothed using a Gaussian filter. Local maxima of the distance map were detected and used as markers for watershed segmentation. Marker-controlled watershed was then applied to the inverted distance map to obtain individual nuclear regions. Each segmented region was characterized using standard morphological and intensity features, including area, solidity, eccentricity, and mean anti-NeuN intensity. To exclude segmentation artifacts and nuclear aggregates, regions were filtered based on morphological criteria: solidity > 0.9, eccentricity < 0.85, and area less than twice the median area of all detected objects. To identify neuronal nuclei, an intensity-based threshold was applied. Specifically, Otsu thresholding was performed on the distribution of mean anti-NeuN intensities across all candidate nuclei within an image, and only regions with mean anti-NeuN intensity exceeding this threshold were retained.

We analyzed 8 SZ and 8 HC cases (Supplementary Table S1). In each case 4 tissue sections were made, and 3 images were made for each section. Differences in area between groups aggregated by age and diagnosis were assessed using the Mann–Whitney U test.

##### General chromatin features analysis

To examine how contact probability changes as a function of genomic distance, Hi-C matrices binned at 10 kb resolution were processed using the expected_cis function from the cooltools package [[2]](https://www.zotero.org/google-docs/?8gm9l0). This function calculates the mean intra-chromosomal contact frequency relative to genomic separation. Smoothing and aggregating data across all autosomes were applied. For each Hi-C sample, the resulting profiles were saved for downstream analysis. Samples were then grouped by diagnostic cohort and in aging analysis by donors’ age. For comparison between diagnostic cohorts, the mean expected contact frequency scaling plot, and average cis-trans chromosomal contacts were calculated. Scaling plot was calculated for each genomic distance bin to generate representative scaling curves within each group. These average contact probability decay profiles were normalized to start from the same point, visualized on log–log plots, showing contact probability as a function of genomic distance. Cis-Trans contacts were calculated as average contacts within each chromosome and between each pair of chromosomes. Chromosomes X and Y, as well as mitochondrial sequences, were excluded from all analyses.

For comparison of general chromatin features in aging groups, the ratio of the mean value of each metric in old donors to that in young donors was calculated, referred to as the old-to-young ratio. Additionally we calculated the old-to-young ratio of Interchromosomal Fraction of Interactions (ICF). ICF for each sample was calculated as the average value of interactions with bins from other chromosomes to the average value of total interactions with bin for each genomic bin. For amplitude estimation of polymer scaling, we calculated the ratio of each old sample to each young sample within SZ and HC cohorts.

##### Identification and analysis of compartments

To identify compartments from Hi-C data, firstly we first applied Inspectro [[3]](https://www.zotero.org/google-docs/?Hx0DHX), v0.2.0, to cluster genomic bins based on the eigenvector decomposition. For each individual Hi-C map, the following parameters were used for clusterization: “binsize: 100000”, “n_eigs: 5”, “n_clusters: [16]” and “decomp_mode: 'trans'”; for each merged Hi-C map, the following parameters were used for clusterization: “binsize: 50000”, “n_eigs: 5”, “n_clusters: [16]” and “decomp_mode: 'trans'”. Next, for each chromosome arm we selected 5 principal components of the Hi-C matrix that correlated the most with genomic GC content using the eigs_cis() function from the cooltools package with parameters: “n_eigs=5”, “sort_metric='spearmanr'”. Among the resulting compartment tracks derived from eigenvector decomposition, we retained the one showing the highest Pearson correlation coefficient with the cooltools principal components. To get the resulting compartment tracks for each chromosomal arm, we selected the principal component that showed the highest Pearson correlation coefficient with genomic bins clusterisation from Inspectro. In some cases, there were 2 or more vectors that had a close correlation, among which the most suitable one was selected by visual inspection. Chromosomes Y and M and short chromosomal arms: chr13p, chr14p, chr15p, chr21p, chr22p - were ignored. The limma package (Ritchie et al., 2015) was used to identify bins exhibiting differences in compartment tracks between HC and SZ cohorts. The code for saddle plots was adapted from the cooltools tutorial [[4]](https://www.zotero.org/google-docs/?Pfc8QX). To calculate compartment interaction strength, the average intensity of the top 20% of corresponding interactions derived from the saddle matrix was used.

##### Chromatin loops processing method

To reconcile minor positional discrepancies in loop coordinates across samples, loops falling within spatial proximity were considered equivalent using a DBSCAN-based grouping strategy (scikit-learn), where co-clustered loops were treated as a single consensus interaction. To mitigate potential false negatives inherent to the cooltools loop-calling procedure described in the corresponding section, a supplementary customized loop search was performed. In cases where no loops were initially identified within a cluster for a given sample, we examined regions with the highest signal intensity relative to surrounding areas within a given cluster. Regions with mean intensity values consistent with called chromatin loops in that specific dataset were reclassified as loops and incorporated into the final analysis. Resulting clusters were then subjected to sequential quality filters, removing those with insufficient contributing samples or below-threshold signal intensities. The final clusters were used in the following analyses. The intersection of the loop anchors with other features presented in .bed or .bedpe files was performed with pybedtools package.

Differential chromatin loop analysis was focused on identifying upregulated and downregulated loops between SZ and HC groups. An approach involving statistical assessments of loop intensities was employed. First, median loop intensity values from each group were calculated for loops within one cluster. Subsequently, the Mann-Whitney U test was applied to evaluate significant intensity changes.

##### Variant calling from Hi-C data and detection of variant-sensitive interactions

Variant calling from Hi-C reads aligned with distiller-nf was performed using the FreeBayes package [[5]](https://www.zotero.org/google-docs/?VcxupA), with indel observations removed from the input. Variants were retained if they met the following criteria: the fraction and number of observations supporting an alternate allele within a single individual were at least 0.33 and 10, respectively; the minimum coverage was at least 10; and the base quality was at least 20. Only SNP positions from SNPdb (build 151) were considered. In addition, to obtain high-confidence variants, we retained only those covered by at least 30 reads in any sample, with mapping quality > 50 and QUAL > 40. This approach yielded 496,779 positions with variants, 468,265 of which were variable across samples. To assess the reliability of the variant calls, we compared them with genotyping-derived variants at matched positions on chromosome 18. The concordance between variants identified from genotyping and Hi-C data was 99.75%.

Overall, this analysis was designed to focus on contacts generated specifically by the genomic position carrying the SNP, rather than on conventional bin-level contacts, where the signal may also arise from any position within the bin. For this purpose, read pairs with one read overlapping a SNP position were extracted for each sample. Only read pairs with distance of less than 1 Mb were retained for downstream analysis. Reads paired to those overlapping SNP positions were aggregated by distance from the SNP into 100-kb bins, resulting in 10 bins upstream and 10 bins downstream of each SNP. Data from all samples were then aggregated by variants, producing a table containing the variant, the number of interactions between the SNP position and each 100-kb bin, the distance from the SNP to the bin, and the total number of reads covering the SNP. We also included GC content, mappability, fragment size, expression in the interacting bin, and the difference in PC1 values between the interacting regions. This table was used as input to a generalized linear model with Poisson distribution, which was used to fit the data distribution and estimate interaction probabilities for calculation of P values. P values were adjusted using the Benjamini–Hochberg correction, and a standard threshold of 0.05 was used to define significant contacts. An interaction was defined as allele-specific if it was significant for one variant but not for another. In the last case, we imposed an additional threshold on the q-value of > 0.1. We excluded approximately 8.2% of significant contacts for which all three variant classes were represented among samples (two homozygous and one heterozygous variant), focusing primarily on comparisons between heterozygous and homozygous variants. This analysis identified 251 variant-sensitive significant interactions among 9716 contacts formed by SNP positions and 100-kb bins within 1 Mb. We further excluded interactions occurring within the 100-kb bin containing the SNP, retaining 103 interactions for downstream analysis. The model was built using the discrete_model module of the statsmodels package [[6]](https://www.zotero.org/google-docs/?4IrnbT).

The difference of averaged heatmaps centred on significant variant-sensitive contacts was plotted as follows: for each contact, Hi-C maps from samples were averaged across variants, generating two mean Hi-C map fragments centred on the interaction. The difference between these maps was then calculated, resulting in a set of heatmaps characterized by enrichment or depletion of the central contact. These heatmaps were averaged across all contacts, with weights of +1 for enrichments and −1 for depletions.

The annotation of interacting bins or SNP positions was based on the intersection with coordinates of genes (full length). Primary eQTL–gene linkages were retrieved from the MetaBrain project (european population, cortical data) [[7]](https://www.zotero.org/google-docs/?eViBWe). Then, we focused on interactions involving intergenic SNPs and bins containing genes (29 items). For each such gene, we extracted the positions of its primary eQTLs and calculated the minimum distance between the eQTL and the SNP involved in the allele-specific contact over all eQTLs of genes in the interacting bin. The mean distance within all contacts was then compared to means obtained from permutation analysis. Control sets for permutation analysis were generated by randomly sampling from non-significant contacts involving intergenic SNPs paired with gene-containing bins, while preserving the number and between-anchor distance distribution of the initial set of 29 significant contacts . Subsequent steps mirrored the analysis of the observed data, yielding 1000 mean minimal eQTL–SNP distances for comparison.

To assess the transcription of genes interacting with SNP-carrying loci, we focused on genes involved in contacts between regions containing two different genes, within the same gene, or between loci with SNPs in intergenic regions and genes. The control gene set was constructed from SNP-associated interactions that were tested but did not show significant allele-specific effects (i.e., those with an adjusted p-value > 0.05, or interactions that were significant but showed similar effects for both variants). All genes intersecting bins that interact with SNP positions were included in this control set.

##### Age Score construction

Publicly available RNA-seq data on sorted human brain nuclei [[8, 9]](https://www.zotero.org/google-docs/?d5X9QF) were used to construct and evaluate Age Score metrics over diagnosis. Gene expression data were normalized with pyDESeq2 using variance stabilizing transformation (VST). To identify age-associated genes, we first computed Pearson correlation coefficients between gene expression levels and chronological age across samples, selecting genes with an absolute correlation coefficient greater than 0.3. These candidate genes were further analyzed using linear ordinary (for Mendizabal’s dataset) or mixed-effects (for Zhu’s dataset) models implemented with the following specification: Gene Expression ~ Age + Status, schizophrenia treatment was included as a random intercept in Zhu’s dataset model. Genes exhibiting a significant effect of age (q-value < 0.1) after FDR correction were retained for downstream analysis.

To construct the Age Score, expression values of the selected genes were first scaled to reduce the influence of extreme values. For each gene g and sample i, normalized expression xg, i was computed by dividing raw VST expression xg, i by the gene-specific 95th percentile, followed by scaling with the median absolute deviation.

The final Age Score for the sample i was defined as:

${Age Score}_{i}=\frac{1}{|G^{+}|}\sum_{g\in G^{+}} \hat{x}_{g, i}-\frac{1}{|G^{-}|}\sum_{g\in G^{-}} \hat{x}_{g, i}$

where G+ and G- denote the sets of genes with positive and negative age-associated regression coefficients, respectively. An ordinary linear model was introduced to analyze Age Score distribution, the next model design was used: Age Score ~ Age * Diagnosis.

### SUPPLEMENTARY FIGURES

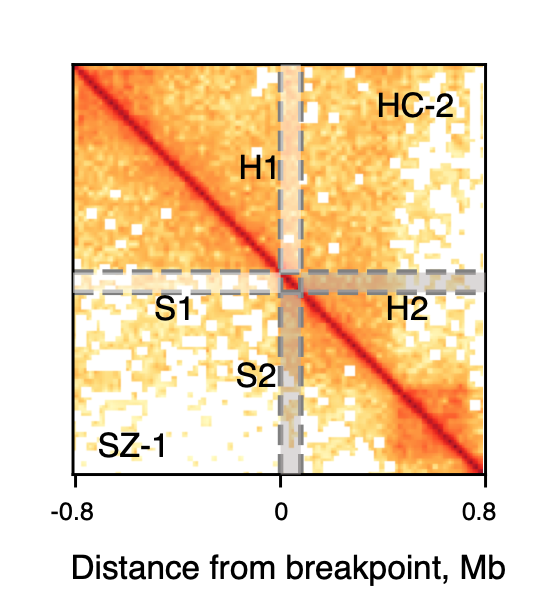

###### Supplementary Figure S1. Schematic representation of the extrusion track intensity calculation: contact frequencies within the white region (S1 for the sample with inversion and H1 for the control) were normalized by the mean interaction frequency within the corresponding grey regions (S2 and H2).

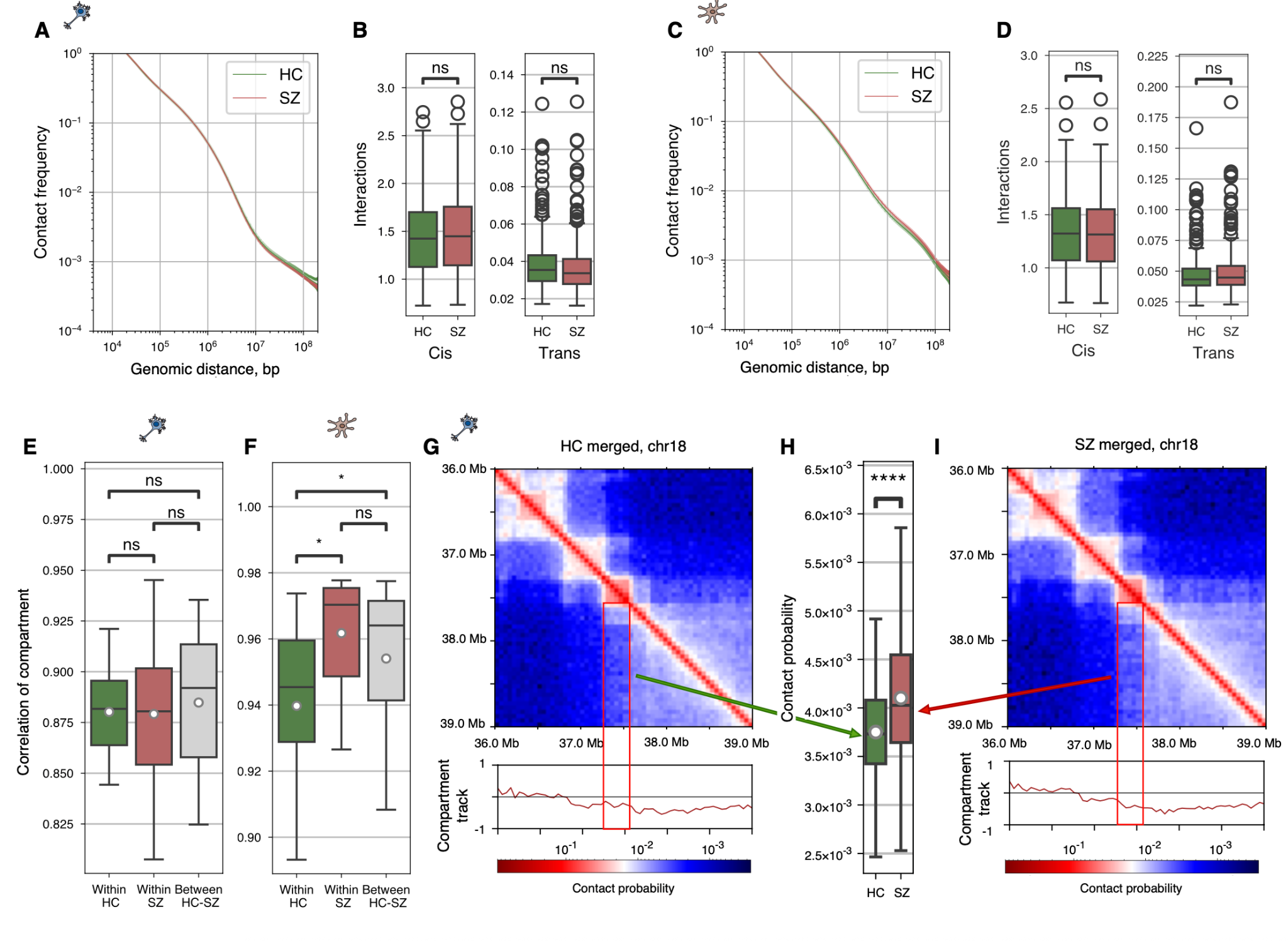

###### Supplementary Figure S2. Analysis of large-scale chromatin features. A, C. HC (green) and SZ (red) average interaction frequencies at various genomic distances in neurons (A) and non-neurons (C). B, D. HC (green) and SZ (red) average interactions within all chromosomes (left boxplots) and between all pairs of chromosomes (right boxplots) in neurons (B) and non-neurons (D). E-F. Correlations of compartments’ eigenvectors within cohorts and between them in neurons (E) (Mann-Whitney U test, comparison between “Within HC” and “Within SZ”: p-value = 0.836; comparison between “Within HC” and “Between HC-SZ”: p-value = 0.502; comparison between “Within SZ” and “Between HC-SZ”: p-value = 0.6125) and non-neurons (F) (Mann-Whitney U test, comparison between “Within HC” and “Within SZ”: p-value = 0.01; comparison between “Within HC” and “Between HC-SZ”: p-value = 0.036; comparison between “Within SZ” and “Between HC-SZ”: p-value = 0.163). G-I. Hi-C maps and compartments profiles near the changed region in merged HC map (G) and SZ map (I) in neurons; contact intensities (H) in the red square region containing the region with changed compartment track for or HC merged map (green) and SZ merged map (red).

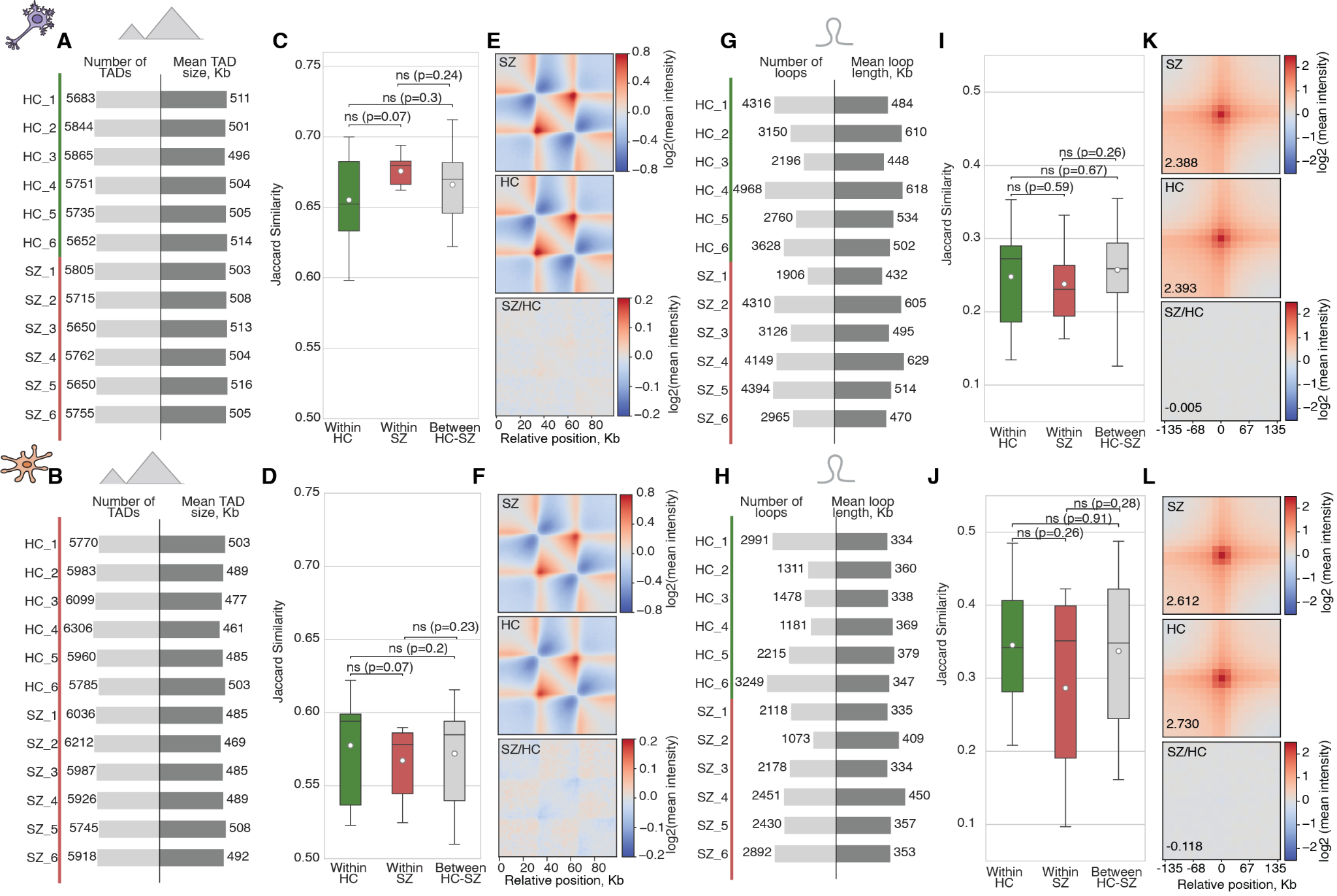

###### Supplementary Figure S3. Fine-scale chromatin features in neurons and non-neuronal cells. A-B. Number of identified TAD borders and average TAD size in neurons (A) and non-neurons (B). C-D. Jaccard similarity for TAD border positions in neurons (C) and non-neurons (D). E-F. Average TAD intensities in schizophrenia (SZ) and healthy (HC) samples, and their ratio, for neurons (E) and non-neurons (F). G-H. Number of identified loops in neurons (G) and non-neurons (H). I-J. Jaccard similarity of loops anchors positions in neurons (I) and non-neurons (J). K-L. Average loop intensities in SZ and HC samples, and their ratio, for neurons (K) and non-neurons (L).

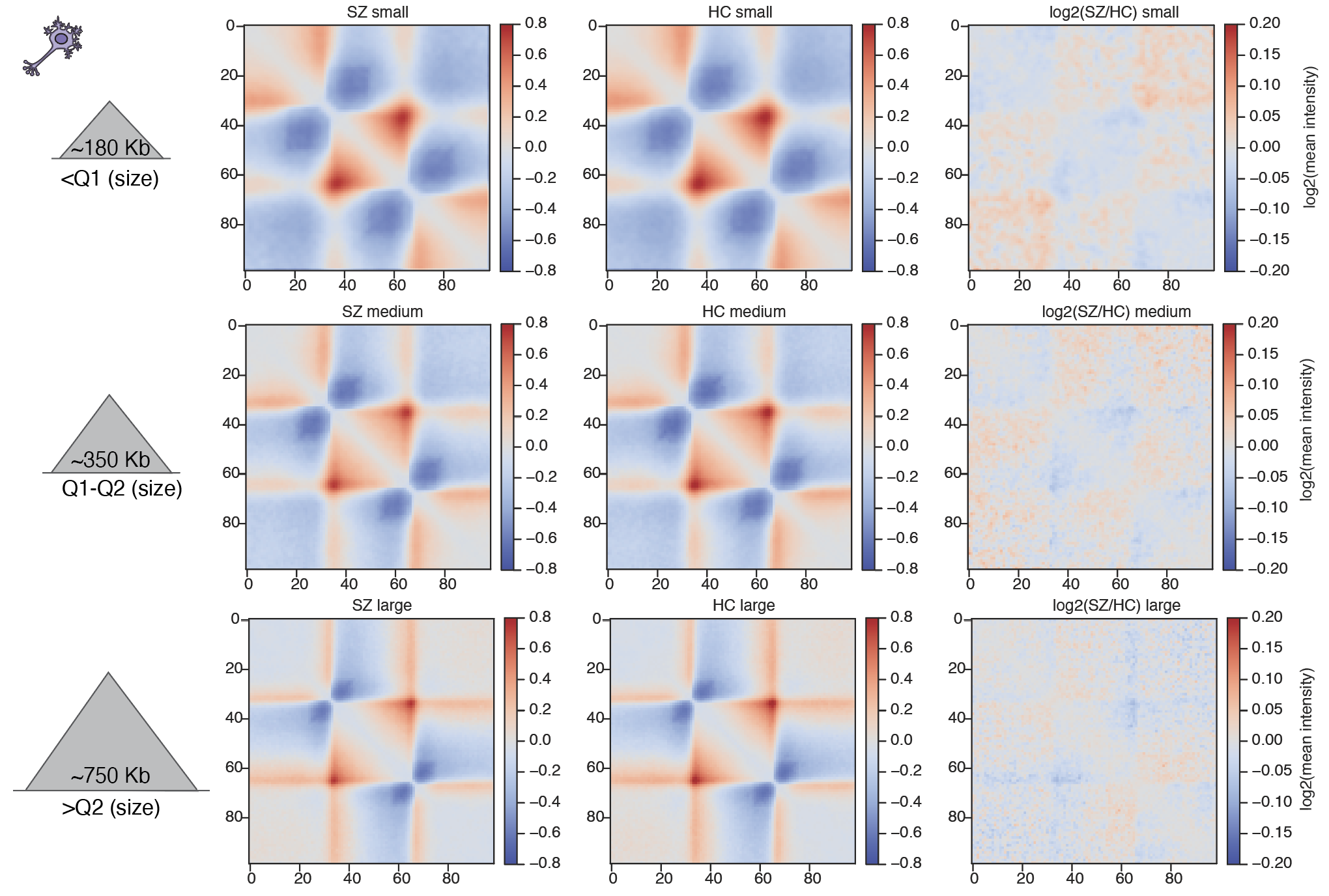

###### Supplementary Figure S4. Average TAD intensities in SZ and HC samples, and their ratio for TADs of different size groups for neurons.

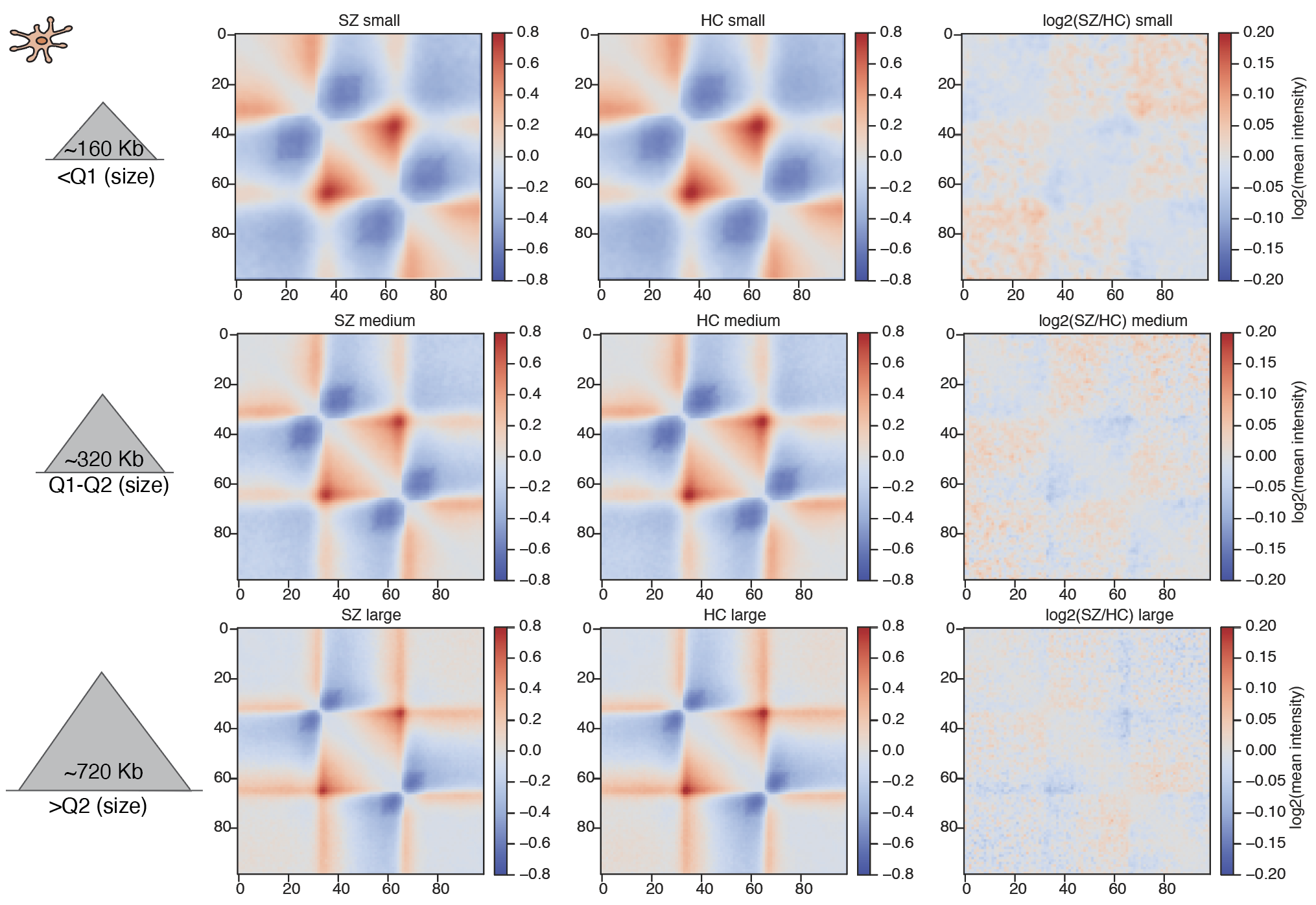

###### Supplementary Figure S5. Average TAD intensities in SZ and HC samples, and their ratio for TADs of different size groups for non-neurons.

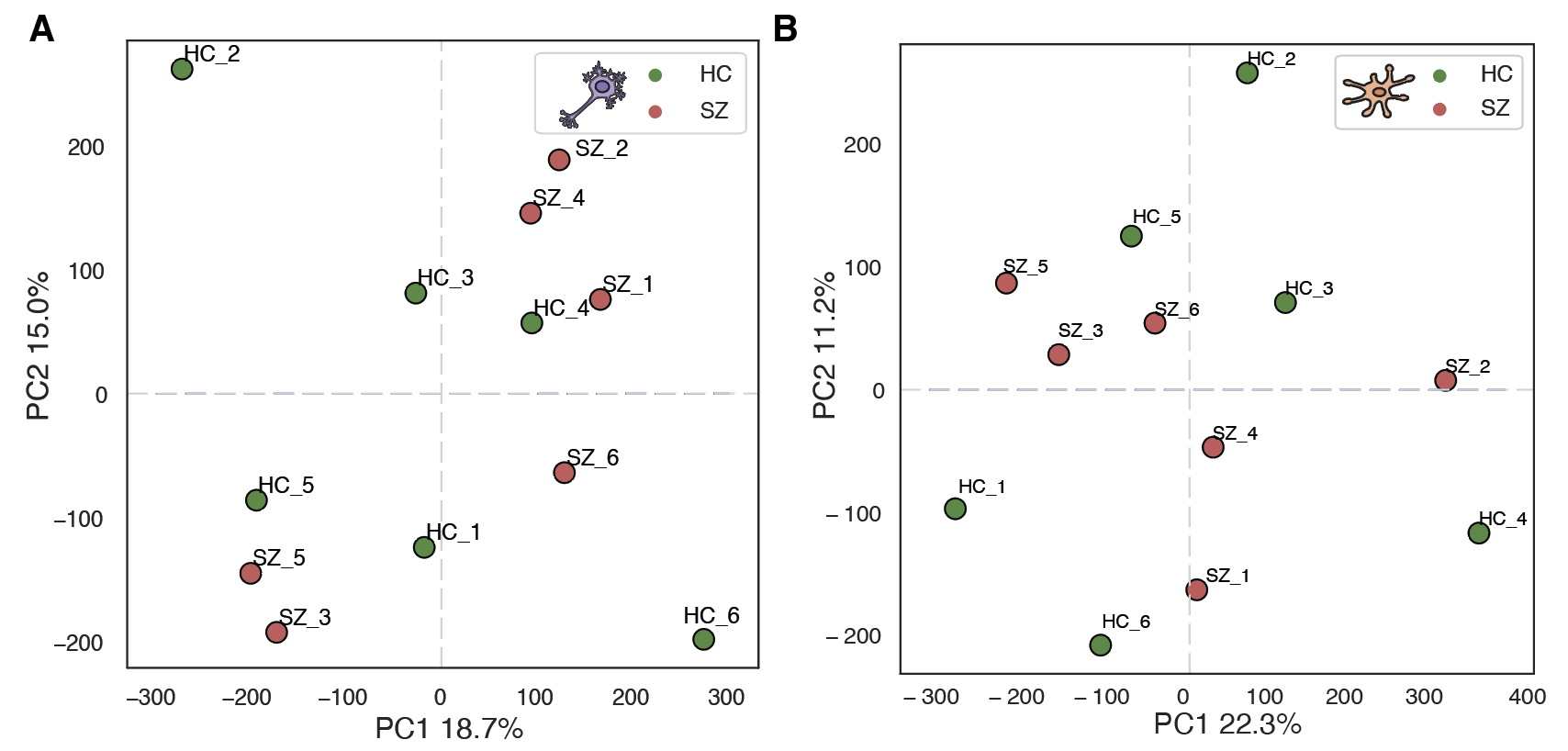

###### Supplementary Figure S6. PCA of insulation score (IS) across analyzed samples for neurons (A) and non-neurons (B).

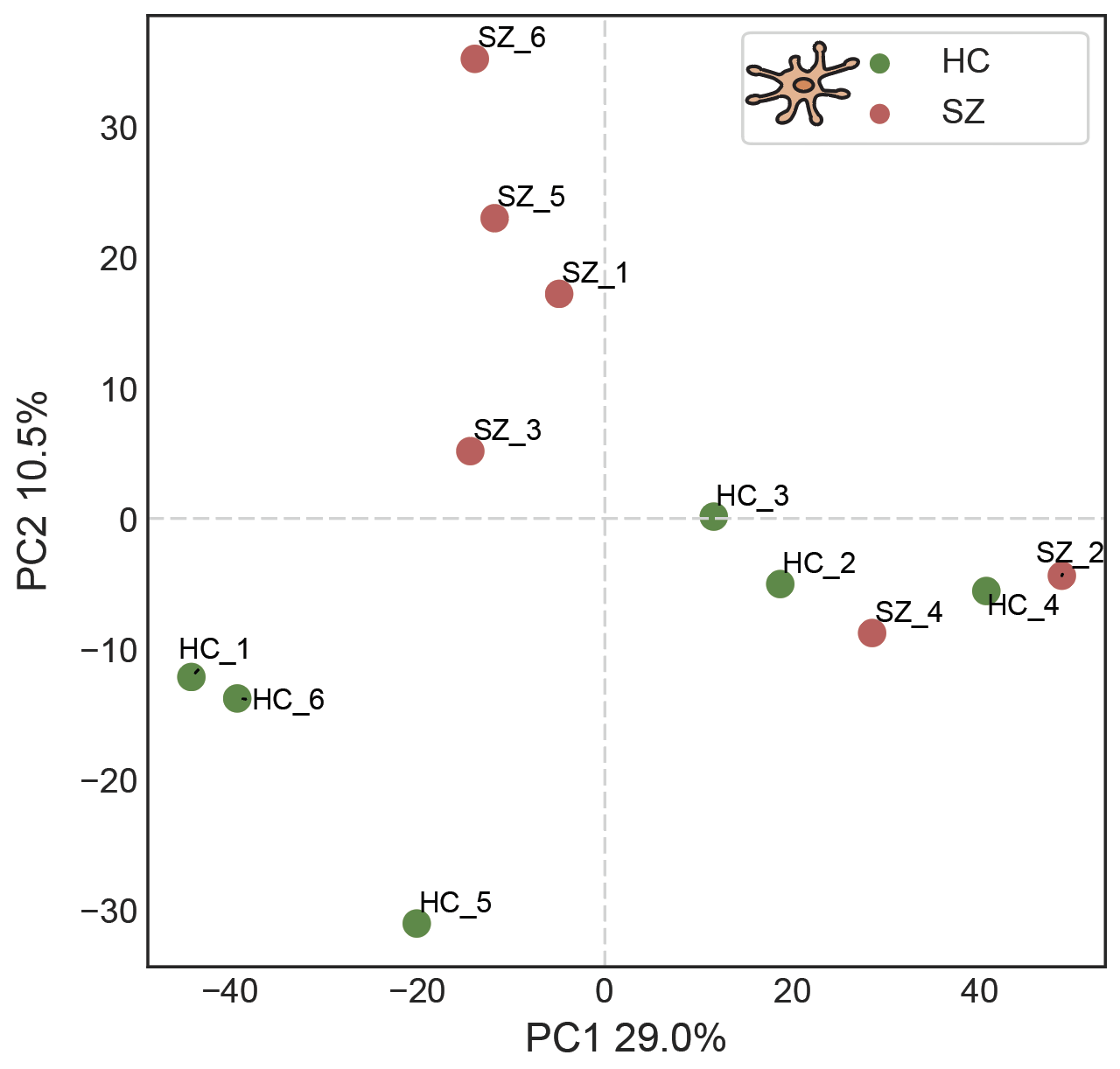

###### Supplementary Figure S7. PCA of loop intensities across analyzed samples for non-neurons.

### SUPPLEMENTARY TABLES

Supplementary Table S1. **Sample metadata** (provided as an additional .xlsx file).

Supplementary Table S2. **Significant variant-sensitive interactions** (provided as an additional .xlsx file).
